# Imbalanced amplification: A mechanism of amplification and suppression from local imbalance of excitation and inhibition in cortical circuits

**DOI:** 10.1101/201269

**Authors:** Christopher Ebsch, Robert Rosenbaum

## Abstract

Understanding the relationship between external stimuli and the spiking activity of cortical populations is a central problem in neuroscience. Dense recurrent connectivity in local cortical circuits can lead to counterintuitive response properties, raising the question of whether there are simple arithmetical rules for relating circuits’ connectivity structure to their response properties. One such arithmetic is provided by the mean field theory of balanced networks, which is derived in a limit where excitatory and inhibitory synaptic currents precisely balance on average. However, balanced network theory is not applicable to some biologically relevant connectivity structures. We show that cortical circuits with such structure are susceptible to an amplification mechanism arising when excitatory-inhibitory balance is broken at the level of local subpopulations, but maintained at a global level. This amplification, which can be quantified by a linear correction to the classical mean field theory of balanced networks, explains several response properties observed in cortical recordings.

## 1 Introduction

Information about a sensory stimulus is passed along a hierarchy of neural populations, from subcortical areas to the cerebral cortex where it propagates through multiple cortical areas and layers. Within each layer, lateral synaptic connectivity shapes the response to synaptic input from upstream layers and populations. In a similar manner, lateral connectivity shapes the response of cortical populations to artificial, optogenetic stimuli. The densely recurrent structure of local cortical circuits can lead to counter-intuitive response properties [57, 41, 2, 43, 10], making it difficult to predict or interpret a population’s response to natural or artificial stimuli. This raises the question of whether there are underlying arithmetic principles through which one can understand the relationship between a local circuit’s connectivity structure and its response properties.

In principle this relationship could be deduced from detailed computer simulations of the neurons and synapses that comprise the circuit. In practice, such detailed simulations can be computationally expensive, depend on a large number of unconstrained physiological parameters, and their complexity can make it difficult to pinpoint mechanisms underlying observed phenomena. In many cases, however, one is not interested in how the response of each neuron is related to the detailed connectivity between every pair of neurons. Relevant questions are often more macroscopic in nature, *e.g.* “How does increased excitation to population A affect the average firing rate of neurons in population B?” For such questions, it is sufficient to establish a relationship between macroscopic connectivity structure and macroscopic response properties.

One such approach is provided by the mean-field theory of balanced networks [58, 59, 48, 45, 30], which is derived in the limit of a large number of neurons and a resulting precise balance of strong excitation with strong inhibition. This notion of precise balance implies a simple relationship between the macroscopic structure of connectivity and firing rates, and balanced network models naturally produce the asynchronous, irregular spiking activity that is characteristic of cortical recordings [58, 59, 47, 49]. However, classical balanced network theory has some critical limitations. While cortical circuits do appear to balance excitation with inhibition, this balance is not always as precise and spike trains are not as asynchronous as the theory predicts [20, 40, 11, 12, 37, 14, 16]. Moreover, precise balance is mathematically impossible under some biologically relevant connectivity structures [48, 45, 30], implying that the classical theory of balanced networks is limited in its ability to model the complexity of real cortical circuits.

We show that cortical circuits with structure that is incompatible with balance are susceptible to an amplification mechanism arising when excitatory-inhibitory balance is broken at the level of local subpopulations, but maintained at a global level. This mechanism of “imbalanced amplification” can be quantified by a linear, finite-size correction to the classical mean field theory of balanced networks that accounts for imperfect balance and local imbalance. Through several examples, we show that imbalanced amplification explains several experimentally observed cortical responses to natural and artificial stimuli.

## 2 Results

### 2.1 The arithmetic of imprecise balance in cortical circuits

We begin by reviewing and demonstrating the classical mean-field theory of balanced networks and a linear correction to the large network limit that the theory depends on. A typical cortical neuron receives synaptic projections from thousands of neurons in other cortical layers, cortical areas or thalamus. These long range projections are largely excitatory and provide enough excitation for the postsynaptic neuron to spike at a much higher rate than the sparse spiking typically observed in cortex. The notion that excitation to cortical populations can be excessively strong has been posed in numerous studies and is typically resolved by accounting for local, lateral synaptic input that is net-inhibitory and partially cancels the strong, net-excitatory external synaptic input [19, 53, 58, 3, 51, 41]. Balanced network theory takes this cancellation to its extreme by considering the limit of large external, feedforward synaptic input that is canceled by similarly large local, recurrent synaptic input. In this limit, a linear mean-field analysis determines population-averaged firing rates in terms of the macroscopic connectivity structure of the network [58, 59].

To demonstrate these notions, we first simulated a recurrent network of *N*_*E*_ = 4000 excitatory (population *E*) and *N*_*I*_ = 1000 inhibitory spiking neurons (population *I*) receiving synaptic connections from an “external” population (*X*) of *N*_*X*_ = 4000 excitatory neurons modeled as Poisson processes. Cortical circuits are often probed using optogenetic methods to stimulate or suppress targeted neuronal sub-populations [7, 15]. As a simple model of optogenetic stimulation of cortical pyramidal neurons, we added an extra inward current to all neurons in population *E* halfway through the simulation (Fig. 1a). Neurons in the local population (*E* and *I*) were modeled using the adaptive exponential integrate-and-fire (AdEx) model, which accurately captures the responses of real cortical neurons [8, 25, 26]. Connectivity was random with each neuron receiving 800 synaptic inputs on average and postsynaptic potential amplitudes between 0.19 and 1.0 mV in amplitude. The recurrent network produced asynchronous, irregular spiking (Fig. 1b), similar to that observed in cortical recordings [53, 52, 47, 17]. Firing rates in populations *E* and *I* were similar in magnitude to those in population *X* and were increased by optogenetic stimulation (Fig. 1c). As predicted by balanced network theory, local synaptic input (from *E* and *I* combined) was net-inhibitory and approximately canceled the external input from population *X* and artificial stimulation combined (Fig. 1d).

**Figure 1:**
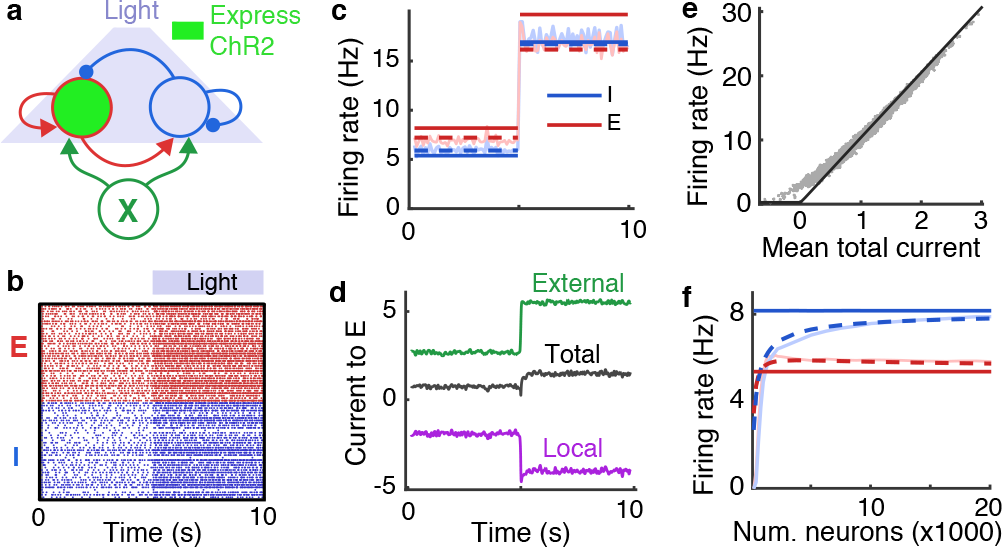
Imprecise balance under optogenetic stimulation. **a)** Schematic. A population of recurrently connected excitatory (red) and inhibitory (blue) spiking neuron models receive synaptic input from an external population (*X*; green) of Poisson-spiking neurons. Optogenetic stimulation of excitatory neurons was modeled by an extra inward current to the excitatory population at 5s. **b)** Spike rasters from 50 randomly selected excitatory (red) and inhibitory (blue) neurons from recurrent network. **c)** Average firing rate of excitatory (red) and inhibitory (blue) neurons in the recurrent network from simualtions (light solid), from the balanced network approximation (Eq. (3); solid dark) and from the corrected approximation (Eq. (4); dashed). **d)** Mean synaptic currents to 200 randomly selected excitatory neurons in the recurrent network from external inputs (*X*; green), from the local population (*E* + *I*; purple) and the total synaptic current (black). Currents are measured in units of the neurons’ rheobase here and elsewhere (rheobase/*C*_*m*_=10.5 V/s). **e)** Mean firing rates plotted against mean input currents to all neurons in populations *E* and *I* (gray dots) and a rectified linear fit to their relationship (black line). **f)** Mean firing rates from identical simulations without stimulation except the total number of neurons, *N*, in the recurrent network was modulated while scaling synaptic weights and connection probabilities so that 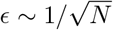 (see Methods). Solid light curves are from simulations, solid dark from Eq. (3), and dashed from Eq. (4).

#### 2.1.1 A review of the mean field theory of balanced networks

To capture the notion that the net external synaptic input to neurons is strong, we define the small number,

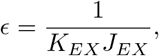

where *K*_*EX*_ = *p*_*EX*_ *N*_*X*_ is the average number of external synaptic projections received by each neuron in *E* from all neurons in *X*, *p*_*EX*_ is connection probability, and *J*_*EX*_ is the synaptic strength of each connection. Specifically, *J*_*EX*_ is the total postsynaptic current induced in a postsynaptic neuron in *E* by a single spike in a presynaptic neuron in *X*. Hence, 1/*ϵ* quantifies the synaptic current that would be induced in each neuron in *E* (on average) if *every* neuron in *X* spiked once simultaneously. Using this convention, the mean synaptic input to each neuron in populations *E* and *I* from all sources can be written in vector form as

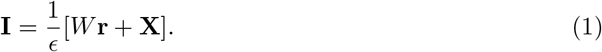

where **I** = [*I*_*E*_ *I*_*I*_]^*T*^ (superscript *T* denotes the transpose) is the vector of mean synaptic input to neurons in each population and similarly for their mean rates, **r** = [*r*_*E*_ *r*_*I*_]^*T*^. The rescaled external synaptic input, **X** = [*X*_*E*_ *X*_*I*_]^*T*^, is given by

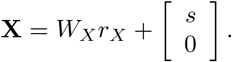

where *r*_*X*_ is the average rate of neurons in population *X* and *s*/*ϵ* is the strength of the inward current induced by optogenetic stimulation (*s* = 0 when stimulation is off). The recurrent and feedforward mean-field connectivity matrices are given by

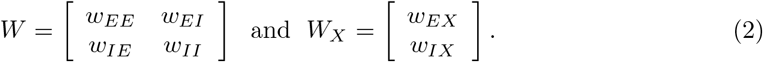

respectively where *w*_*ab*_ = *K*_*ab*_*J*_*ab*_/(*K*_*EX*_*J*_*EX*_) quantifies the relative number, *K*_*ab*_ = *p*_*ab*_*N*_*b*_, and strength, *J*_*ab*_, of synaptic connections from population *b* to *a*. To achieve moderate firing rates when *ϵ* is small, local input, *W* **r**, must be net-inhibitory and partially cancel the strong external excitation, **X**, in Eq. (1).

Balanced network theory [58, 59] takes this cancellation to its extreme by considering the limit of large number of neurons, *N* = *N*_*E*_ + *N*_*I*_, while scaling connection strengths and probabilities in such a way that 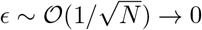. Under this scaling, Eq. (1) would seem to imply that mean synaptic currents diverge in the limit, but this divergence is avoided in balanced networks by a precise cancellation between external and recurrent synaptic input. To achieve this cancellation, firing rates must satisfy the mean-field balance equation,

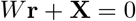

in the large *N* limit, so that [58, 59, 48, 45, 30]

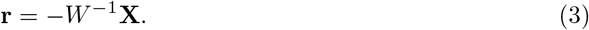

Hence, balanced network theory provides a closed form, linear expression for firing rates in the large network limit. Generally speaking, the firing rate of a neuron depends nonlinearly on the mean and variance of its input current [3, 9, 46]. Notably, however, the derivation of the fixed point in Eq. (3) did not require us to specify the exact form of this dependence. Instead, Eq. (3) represents the unique fixed point firing rates for which synaptic currents remain bounded as *N* → ∞. More specifically, if Eq. (3) is not satisfied as *N* → ∞ then ‖**I**‖ → ∞ (where ‖ · ‖ is the Euclidean norm). The existence of this fixed point does not guarantee that it is stable. Precise, general conditions on the accuracy of Eq. (3) for spiking network models are not known and the investigation of such conditions is outside the scope of this study. However, the approximation tends to be accurate in the *N* → ∞ limit whenever all eigenvalues of *W* have negative real part, the solution in Eq. (3) is strictly positive, and inhibitory synaptic kinetics are sufficiently fast [58, 59, 46, 32, 48, 45, 30]. Indeed, Eq. (3) provides a reasonable, but imperfect approximation to firing rates in our spiking network simulation (Fig. 1c, compare light and dark solid).

Balanced network theory has some critical limitations. Local cortical circuits are, of course, finite in size so the *N* → ∞ (equivalently *ϵ* → 0) limit may not be justified. Moreover, excitation and inhibition in cortex may not be as perfectly balanced and spike trains not as asynchronous as predicted by balanced network theory [20, 40, 11, 12, 55, 37, 14, 16]. More importantly, under many biologically relevant connectivity structures, precise cancellation cannot be realized so Eq. (3) cannot even be applied [48, 45, 30]. We next review a simple, linear correction to Eq. (4) that partially resolves these issues.

#### 2.1.2 A linear correction to precise balance

A correction to Eq. (3) can be obtained by considering *ϵ* non-zero, but this requires making assumptions on the relationship between neurons’ input statistics and firing rates. A simple approximation is obtained by assuming that population-averaged firing rates, **r**, depend only on population-averaged mean inputs, **I**, yielding the fixed points problem **r** = *f*(**I**) = *f*([*W***r** + **X**]/*ϵ*) where *f* is the population-level f-I curve. When *f* is an increasing function over relevant ranges of **I**, this fixed point equation can be re-written as

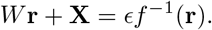

Hence, in strongly coupled networks (*ϵ* small), the shape of f-I curves has a small effect on steady-state firing rates under such an approximation. Indeed, in the *ϵ* → 0 limit, the f-I curve has no effect and firing rates are determined by Eq. (3). This conclusion easily generalizes to the case where *f* also depends on the average temporal variance of neurons’ inputs.

A simple case of this approximation is obtained by using a rectified linear approximation, **r** = *g*[**I**]_+_ where [·]_+_ denotes the positive part. We fit such a function to the relationship between neurons’ mean synaptic inputs and firing rates from our spiking network simulation (Fig. 1e). Assuming that the average firing rates of all populations are positive, this rectified linear approximation produces a linear rate model [13] with mean firing rates given by solving *W***r** + **X** = *ϵ*/*g* **r** to obtain

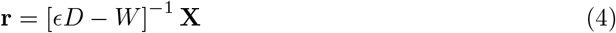

where

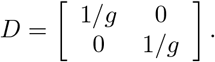

The AdEx neuron model used in our simulations has a nonlinear f-I curve (Fig. 1e; gray dots) and its firing rate depends on all statistics of its input, not just the mean [8, 21]. Nevertheless, the linear approximation in Eq. (4) was accurate in predicting firing rates in our simulations (Fig. 1c, solid), outperforming the balanced network approximation from Eq. (3). This can be explained by the fact that the balanced approximation in Eq. (3) is already somewhat accurate and the linear approximation in Eq. (4) corrects for some of the error introduced by imperfect balance, even though the true dependence of **r** on **I** is nonlinear.

To further investigate the relative accuracy of Eqs. (3) and (4), we repeated the spiking network simulations from Fig. 1a-d while proportionally scaling the number of neurons (*N*_*E*_, *N*_*I*_, and *N*_*X*_) in each population and scaling connection weights and probabilities in such a way that 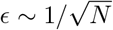 (see Methods). As predicted by balanced network theory, excitatory and inhibitory firing rates increased toward the limit in Eq. (3) (Fig. 1f, compare light and dark solid lines). The linear correction in Eq. (4) tracks this increase in firing rates and is more accurate than the approximation in Eq. (3), particularly for smaller *N* (Fig. 1f, dashed). It is worth noting that, in applying Eq. (4) to obtain the dashed curve in Fig. 1f, we fixed the value of *g* to the one obtained from the simulation in Figs. 1a-e. Hence, a single estimate of the gain yields an accurate approximation even under different parameter values.

The predictive power of Eq. (4) in these examples is, of course, limited by the fact that it was only applied after estimating the gain of the neurons using firing rates obtained in simulations. Moreover, highly nonlinear f-I curves could introduce additional error. However, the purpose of Eq. (4) in this work is to provide a first-order approximation to and general understanding of firing rates in networks under which Eq. (3) cannot be applied. For these purposes, Eq. (4) is sufficient.

### 2.2 Imbalanced amplification under partial optogenetic stimulation

We next show that a more realistic model of optogenetic stimulation breaks the classical balanced state, providing a demonstrative and experimentally relevant example of imbalanced amplification and suppression that explains phenomena observed in recordings from mouse somatosensory cortex.

#### 2.2.1 Firing rates are increased by stimulating fewer neurons

The model of optogenetic stimulation considered in Fig. 1 is somewhat inaccurate since optogenetic stimulation of excitatory neurons is often incomplete. For example, only a fraction of cortical pyramidal neurons express the channelrhodopsin 2 (ChR2) protein targeted in many optogenetic experiments [7, 42, 44, 2]. To more accurately model optogenetic stimulation, we modified the example above so the extra inward current was provided to only 20% of the excitatory neurons (Fig. 2a), modeling ChR2-expressing pyramidal cells. This change produced surprising results. The ChR2-expressing neurons increased their firing rates by a larger amount than they did when all excitatory neurons received the current (Fig. 2b,c; compare to Fig. 1b,c). Hence, counterintuitively, stimulating fewer neurons actually *amplifies* the effects of stimulation on the targeted cells. In contrast, non-expressing excitatory neurons were *suppressed* during stimulation and inhibitory neurons increased their rates, but by a smaller amount than they did under complete stimulation (Fig. 2e; compare to Fig. 1c).

**Figure 2:**
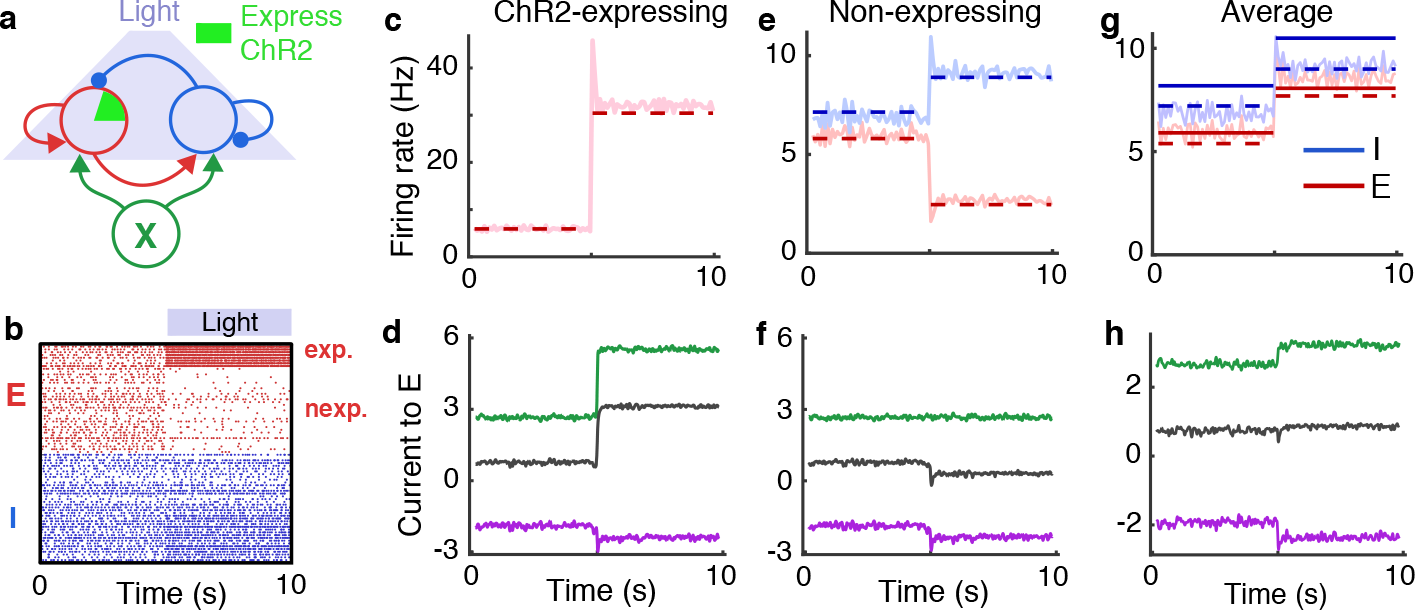
Imbalanced amplification and suppression under partial optogenetic stimulation. Same as Fig. 2 except the inward current was only provided to 20% of the excitatory neurons, modeling ChR2-expressing pyramidal cells. Firing rates of and input current to excitatory neurons are shown separately for ChR2-expressing (c,d) and non-expressing (e,f) neurons, as well as the average over all neurons (g,h). Firing rates predicted by Eq. (3) are not shown in c and e because Eq. (3) is not applicable to those cases.

Similar effects were observed in experiments by Adesnik and Scanziani [2]. In that study, pyramidal neurons in layers (L) 2/3 of mouse somatosensory cortex (S1) were stimulated optogenetically, but only about 23% of the pyramidal neurons expressed ChR2. During stimulation, non-expressing L2/3 pyramidal neurons were suppressed and inhibitory synaptic currents increased, implying an increase in inhibitory neuron firing rates.

To understand these effects, we first extended the mean-field theory above to account for multiple subpopulations by defining

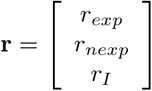

to be the vector of average firing rates for the ChR2-expressing (*exp*), non-expressing (*nexp*) excitatory neurons and inhibitory (*I*) neurons. The vector of average input to the network is again given by Eq. (1) where

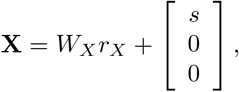

*W*_*X*_ = [*w*_*EX*_ *w*_*EX*_ *w*_*IX*_]^*T*^,

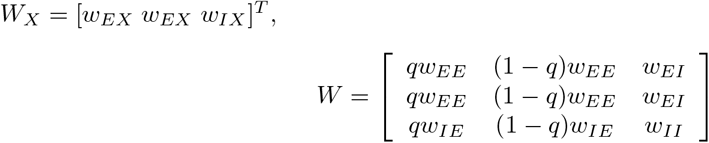

and *q* = 0.2 represents the proportion of neurons that express ChR2.

Importantly, *W* is singular (*i.e.*, not invertible), so classical balanced network theory fails for this example since Eq. (3) cannot be evaluated. More specifically, it is impossible for **I** in Eq. (1) to remain finite as *ϵ* → 0 since there is no vector, **r**, such that *W***r** = −*X*. Intuitively, this can be understood by noting that expressing and non-expressing excitatory neurons receive the same local input on average (Fig. 2d,f, purple), since local connectivity is not specific to ChR2 expression, but they receive different external input during stimulation (Fig. 2d,f, green). Therefore, local synaptic input cannot simultaneously cancel the external input to both subpopulations, so the precise cancellation required by classical balanced network theory cannot be achieved (Fig. 2d,f, black). A similar mechanism has been used to explain a lack of cancellation between positive and negative correlations in balanced networks [60, 49].

#### 2.2.2 Amplification in the nullspace: a general analysis

We now give a general analysis of network responses when *W* is singular. The example of partial optogenetic stimulation is then considered as a special case. If *W* is a singular matrix then only vectors, **X**, that are in the column space of *W* admit solutions to *W***r** + **X** = 0. The column space of *W* is defined as the linear space of all vectors, **u**, such that **u** = *W***r** for some **r**. The column space of a matrix, *W*, is the orthogonal complement of the nullspace of *W*^*T*^. We can therefore decompose

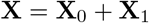

where 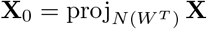 is the projection of **X** onto the nullspace of *W*^*T*^ and **X**_1_ = proj_*C*(*W*_) **X** is the projection onto the column space of *W*. Moreover, note that 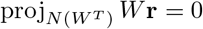 since *W***r** is in the column space of *W*. Therefore, the projection of the total input onto the nullspace of *W*^*T*^ is

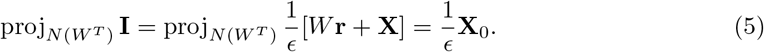

Hence, the projection of the total synaptic input onto the nullspace of *W*^*T*^ is 𝒪(1/*ϵ*) whenever **X** has an 𝒪(1) component in the nullspace of *W*^*T*^. Note that, despite the 1/*ϵ* term in Eq. (1), the total synaptic input, **I**, is 𝒪(1) when balance is realized due to cancellation (as in Fig. 1d). Hence, the singularity of *W* introduces large, 𝒪(1/*ϵ*) synaptic currents where they would not occur if *W* was non-singular. In other words, external input in the nullspace of *W*^*T*^ produces strong synaptic currents in the network. Importantly, this conclusion does not rely on any assumptions about neurons’ f-I curves or other properties. This result is a fundamental property of balanced networks or, more generally, networks receiving strong feedforward input.

To understand the implications of this result on firing rates in the network, however, we must specify an f-I curve. We again consider the linear rate approximation quantified by Eq. (4). Importantly, unlike Eq. (3) from classical balanced network theory, the approximation in Eq. (4) is applicable to this example because it accounts for imperfect cancellation between local and external inputs. Specifically, the regularized matrix, *ϵD* − *W*, is invertible so Eq. (4) can be evaluated even though Eq. (3) cannot. The resulting firing rate solution from Eq. (4) agrees well with spiking network simulations (Fig. 2c,e). Hence, Eq. (4) provides an accurate approximation to firing rates in networks to which classical balanced network theory is not applicable at all.

Eq. (4) also provides a concise mathematical quantification of firing rates when *W* is singular. Namely, if **X**_0_, **X**_1_ ~ 𝒪(1) then firing rates can be expanded as

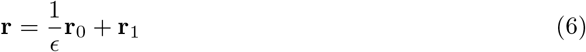

where **r**_0_ is in the nullspace of *W* and **r**_0_, **r**_0_ ~ 𝒪(1). To derive this result, first note that Eq. (4) can be rewritten as

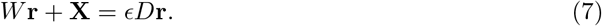

If **X** has components in the nullspace of *W*^*T*^ then we can project both sides of this equation onto this nullspace to obtain

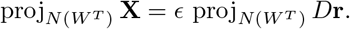

where we again used the fact that 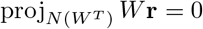 since *W***r** is in the column space of *W*. Since 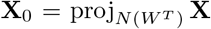 and *D* are assumed 𝒪(1), this equation is only consistent when **r** ~ 𝒪(1/*ϵ*). We can therefore decompose **r** = (1/*ϵ*)**r**_0_ + **r**_1_ where **r**_0_,**r**_1_ ~ 𝒪(1). We next show that **r**_0_ is in the nullspace of *W*. From Eq. (7), we have

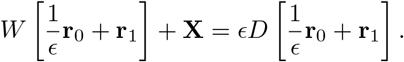

Isolating the 𝒪(1/*ϵ*) terms gives *W***r**_0_ = 0 and therefore **r**_0_ is in the nullspace of *W*. In summary, components of external input in the nullspace of *W*^*T*^ partially break balance to evoke amplified firing rates in the nullspace of *W*.

In the special case that *W* has a one-dimensional nullspace, a more precise characterization of **r**_0_ is possible. Let **v**_0_ be in the nullspace of *W* with ‖**v**_0_‖ = 1. Note that W^*T*^ also has a one-dimensional nullspace (since *W* is a square matrix). Let **v**_*2*_ be in the nullspace of *W*^*T*^ with ‖**v**_2_‖ = 1. Since **r**_0_ is in the nullspace of *W*, we can write **r**_0_ = *a***v**_0_ for some scalar, *a.* Now, dot product both sides of Eq. (7) by **v**_2_ to obtain

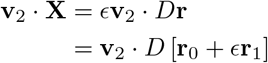

where we have used that **v**_2_ · *W***r** = 0 since **v**_2_ is in the nullspace of *W*^*T*^, which is orthogonal to *W***r** in the column space of *W*. Keeping only 𝒪(1) terms and making the substitution **r**_0_ = *a***v**_0_, we get

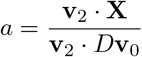

so that

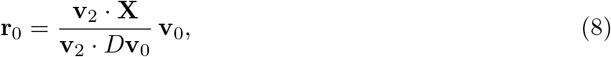

yielding a concise expression for the amplified component of firing rates when *W* has a one-dimensional nullspace. Note that 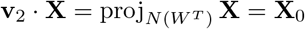 and **v**_0_ is in the nullspace of *W*, so this result is consistent with the more general conclusions above.

#### 2.2.3 Amplification in the nullspace under partial optogenetic stimulation

For the specific example of partial optogenetic stimulation considered in Fig. 2, the nullspace of *W*^*T*^ is spanned by 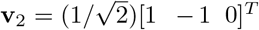 and the projection of **X** onto the nullspace of *W*^*T*^ is **X**_0_ = [*s*/2 −*s*/2 0]^*T*^. The nullspace of *W* is spanned by 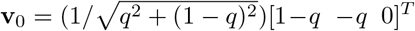. We therefore have **r** = (1/*ϵ*)**r**_0_ + **r**_1_ where Eq. (8) gives

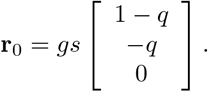

Hence, ChR2-expressing neurons are amplified and non-expressing neurons are suppressed by optogenetic stimulation, as observed in simulations. A more precise result is given by expanding the full approximation from Eq. (4) to obtain

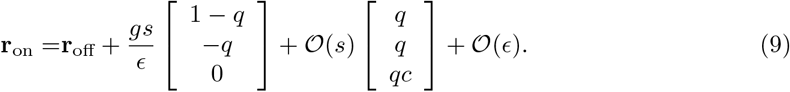

Here, 𝒪(*s*) is a constant proportional to *s*, *c* = |*w*_*IE*_/*w*_*II*_| and **r**_off_ is the vector of firing rates in the balanced, *ϵ* → 0, limit when stimulation is off (*s* = 0). Specifically, **r**_off_ is the unique vector that satisfies *W***r**_off_ + *W*_*X*_*r*_*X*_ = 0, which is solvable even though *W* is singular because W_*X*_*r*_*X*_ is in the column space of *W*, so balance can be maintained when *s* = 0.

The 𝒪(*s*/*ϵ*) term in Eq. (9) quantifies the amplification and suppression observed in simulations: Non-expressing neurons are suppressed by stimulation since −*q* < 0 and the response of ChR2-expressing neurons is amplified since 1 −*q* > 0 and *s*/*ϵ* is large. The 𝒪(*s*) term shows why inhibitory neurons increase their rates by a smaller amount. In summary, the optogenetically induced suppression observed experimentally by Adesnik and Scanziani [2] is a generic feature of balanced or strongly coupled networks under partial stimulation.

#### 2.2.4 Local imbalance with global balance explains intralaminar suppression and interlaminar facilitation

Interestingly, despite the break of balance at the level of ChR2-expressing and non-expressing subpopulations, global balance is maintained in this example. This can be understood by repeating the mean-field analysis above without partitioning neurons into ChR2-expressing and non-expressing sub-populations, thereby quantifying the global average of firing rate of all excitatory neurons. In particular, the average synaptic input, **I** = [*I*_*E*_ *I*_*I*_]^*T*^, to excitatory and inhibitory neurons is given by Eq. (1) where *W* and *W*_*X*_ are as in Eq. (2), and

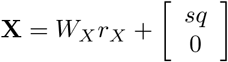

to account for the fact that only a proportion *q* of the excitatory neurons receive the inward current from optogenetic stimulation. In this case, *W* is non-singular so the balanced solution in Eq. (3) is applicable. Indeed, the average firing rates of all excitatory neurons in our spiking network simulation is close to the prediction from Eq. (3) and even closer to the prediction from Eq. (4) (Fig. 2g; compare to Fig. 1c). The average feedforward input to all excitatory neurons is canceled by net-inhibitory local input (Fig. 2h; compare to Fig. 1d). Hence, balance is maintained globally even though the network is imbalanced at the level of ChR2-expressing and non-expressing populations.

In the same study by Adesnik and Scanziani considered above [2], recordings were made in L5, which was not directly stimulated optogenetically, but receives synaptic input from L2/3. Interestingly, despite the fact that most excitatory neurons in L2/3 were suppressed during stimulation, firing rates in L5 increased.

To model these experiments, we interpreted the recurrent network from Fig. 2 as a local neural population in L2/3, which sends synaptic projections to L5 (Fig. 3a). We modeled a neural population in L5 identically to the L2/3 population, except its feedforward input came from excitatory neurons in the L2/3 network, instead of from Poisson-spiking neurons. As in experiments [2], L5 neurons increased their firing rates during stimulation (Fig. 3b) and approximate balance was maintained (Fig. 3c). This can be understood by noting that L5 receives synaptic input sampled from all excitatory neurons in L2/3. Hence, the feedforward excitatory current to L5 neurons increases proportionally to the average excitatory firing rates in L2/3 during stimulation. As we showed above, this average rate increases (Fig. 2e), despite the fact that most excitatory neurons in L2/3 are suppressed by stimulation. Hence, the combination of intralaminar suppression and interlaminar facilitation observed during optogenetic stimulation in experiments [2] results from the fact that the stimulated layer is locally imbalanced, but globally balanced during partial stimulation.

**Figure 3:**
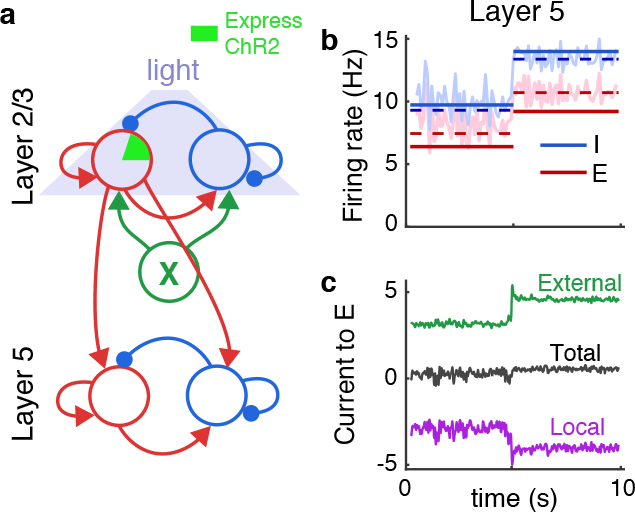
Interlaminar facilitation despite intralaminar suppression under optogenetic stimulation. **a)** Multi-layer network schematic. L2/3 was identical to the recurrent network in Fig. 2 and provided external excitatory input to L5, which had the same internal structure as the L2/3 model. **b)** Excitatory and inhibitory firing rates in L5. **c)** Average synaptic current to randomly sampled excitatory neurons in L5.

#### 2.2.5 Imbalanced amplification of weak stimuli

Sufficiently small *ϵ* or large *s* would introduce negative rates in Eq. (9), representing a regime in which non-expressing neurons cease spiking and the firing rate of ChR2-expressing neurons saturate at a high value. In this sense, firing rates do not truly have a 𝒪(1/*ϵ*) component for *ϵ* very small. However, smaller values of *ϵ* allow weak stimuli (small *s*) to be strongly amplified. Strictly speaking, if one takes *s* ~ 𝒪(*ϵ*), then under the linear approximation in Eq. (4), partial optogenetic stimulation would have an 𝒪(1) effect on the average firing rate of stimulated and unstimulated subpopulations, but an 𝒪(*ϵ*) effect on globally averaged firing rates. In practical terms, this means that, in strongly coupled networks (*ϵ* small), partial optogenetic stimuli can have a moderate effect on the firing rates of stimulated neurons while having a negligible effect on the average firing rates of all excitatory neurons.

To demonstrate this idea, we repeated the simulations from Fig. 2 in a network with four times as many neurons (*N* = 2 × 10^4^) where synaptic weights and probabilities were scaled so that 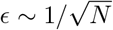 (as in Fig. 1f) and we reduced stimulus strength, *s*, as well. In this simulation, ChR2-expressing neurons’ firing rates nearly doubled (Fig. 4a) and non-expressing neurons were noticeably suppressed (Fig. 4b). However, the change in the average firing rate of all excitatory neurons was nearly imperceptible (Fig. 4c) and similarly for the firing rates of inhibitory neurons (Fig. 4b,c). As a result, the firing rates in a downstream layer were unnoticeably modulated during stimulation (Fig. 4d; compare to Fig. 3). This effect could mask the effects of optogenetic stimulation in recordings.

**Figure 4:**
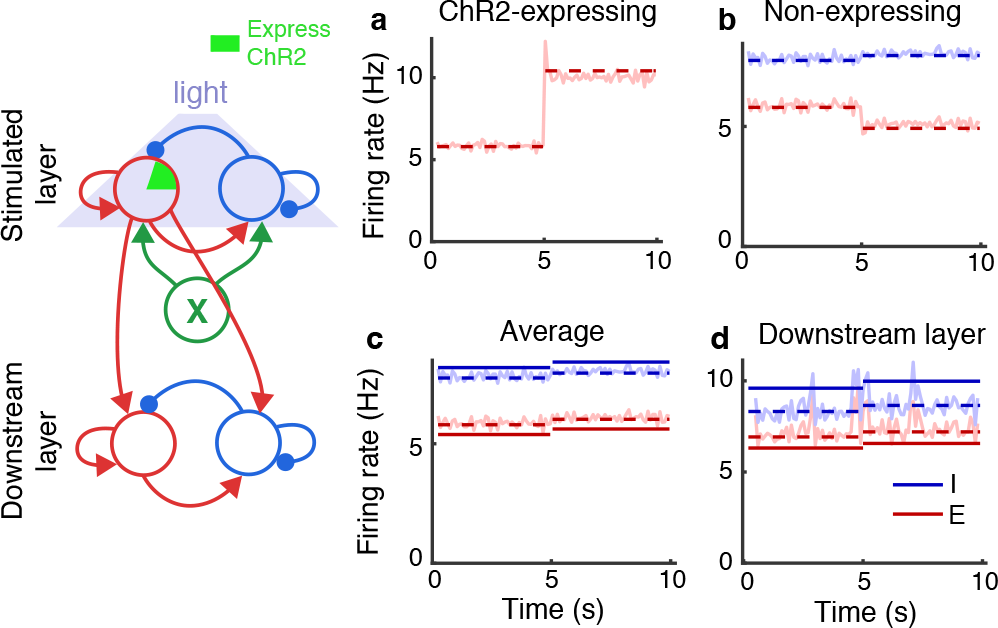
Imbalanced amplification of weak stimuli. **a-c)** Same as Fig. 2b,e,g except with *N* increased by a factor of four, *ϵ* decreased by a factor of two, and a weaker stimulus. **d)** Same as Fig. 3b except using excitatory neurons from the recurrent network in a-c as the feedforward input.

#### 2.2.6 mbalanced amplification with nearly singular connectivity matrices

An apparent limitation of the results above is that they rely on the singularity of the connectivity matrix, *W*. Singularity is a fragile property of matrices that arises from structural symmetries. In the example above, singularity arises from our implicit assumption that local synaptic connectivity is independent of whether neurons express ChR2. Even a slight difference in connectivity to or from ChR2-expressing neurons would make *W* non-singular so that its nullspace would be empty, rendering Eq. (6) vacuous. We now show that Eq. (6) and the surrounding analysis naturally extends to connectivity matrices that are approximately singular, with similar overall conclusions.

A matrix, *W*, is singular if it has *λ* = 0 as an eigenvalue. A matrix can therefore be considered approximately singular if it has an eigenvalue with small magnitude. Specifically, let *λ* be an eigenvalue of *W* with |λ| ≪ 1. Note that *λ* is also an eigenvalue of *W*^*T*^. Now let **v** be the associated eigenvector so that *W*^*T*^v = *λ***v** and assume that ‖**v**‖ = 1 without loss of generality. Take the projection of each term in Eq. (7) onto the subspace spanned by v to get

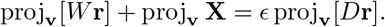

Now note that proj_**v**_[*W***r**] = *λ* proj_**v**_ **r**. Hence,

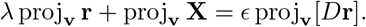

If proj_**v**_ *X* ~ 𝒪(1) and proj_**v**_[*D***r**] ~ proj_**v**_ **r** then this implies

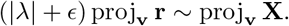

Hence,

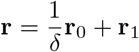

where δ = |*λ*|+*ϵ*. This generalizes Eq. (6) to the case where *W* is only approximately singular. In summary, the mechanism of imbalanced amplification is a general property of strongly coupled networks with singular or nearly singular connection matrices.

We next show that networks with connection probabilities that depend on continuous quantities like distance or tuning preference necessarily have singular or nearly singular connectivity kernels and are therefore naturally susceptible to the amplification and suppression mechanisms described above.

### 2.3 Imbalanced amplification and suppression in continuously indexed networks

So far we considered networks with discrete subpopulations. Connectivity in many cortical circuits depends on continuous quantities like distance in physical or tuning space. To understand how the amplification and suppression mechanisms discussed above extend to such connectivity structures, we next considered a model of a visual cortical circuit. We arranged 2 × 10^5^ AdEx model neurons (80% excitatory and 20% inhibitory) on a square domain, modeling a patch of L2/3 in mouse primary visual cortex (V1). Neurons received external input from a similarly arranged layer of 1.6 × 10^5^ Poisson-spiking neurons, modeling a parallel patch of L4 (Fig. 5a). We additionally assigned a random orientation preference to each neuron, modeling the “salt-and-pepper” distribution of orientation preferences in mouse V1. Connectivity was probabilistic and, as in cortex [23, 28, 33], inter- and intralaminar connections were more numerous between nearby and similarly tuned neurons. Specifically connection probability decayed like a Gaussian as a function of distance in physical and orientation space (Fig. 5b), where distance in both spaces was measured using periodic boundaries.

**Figure 5:**
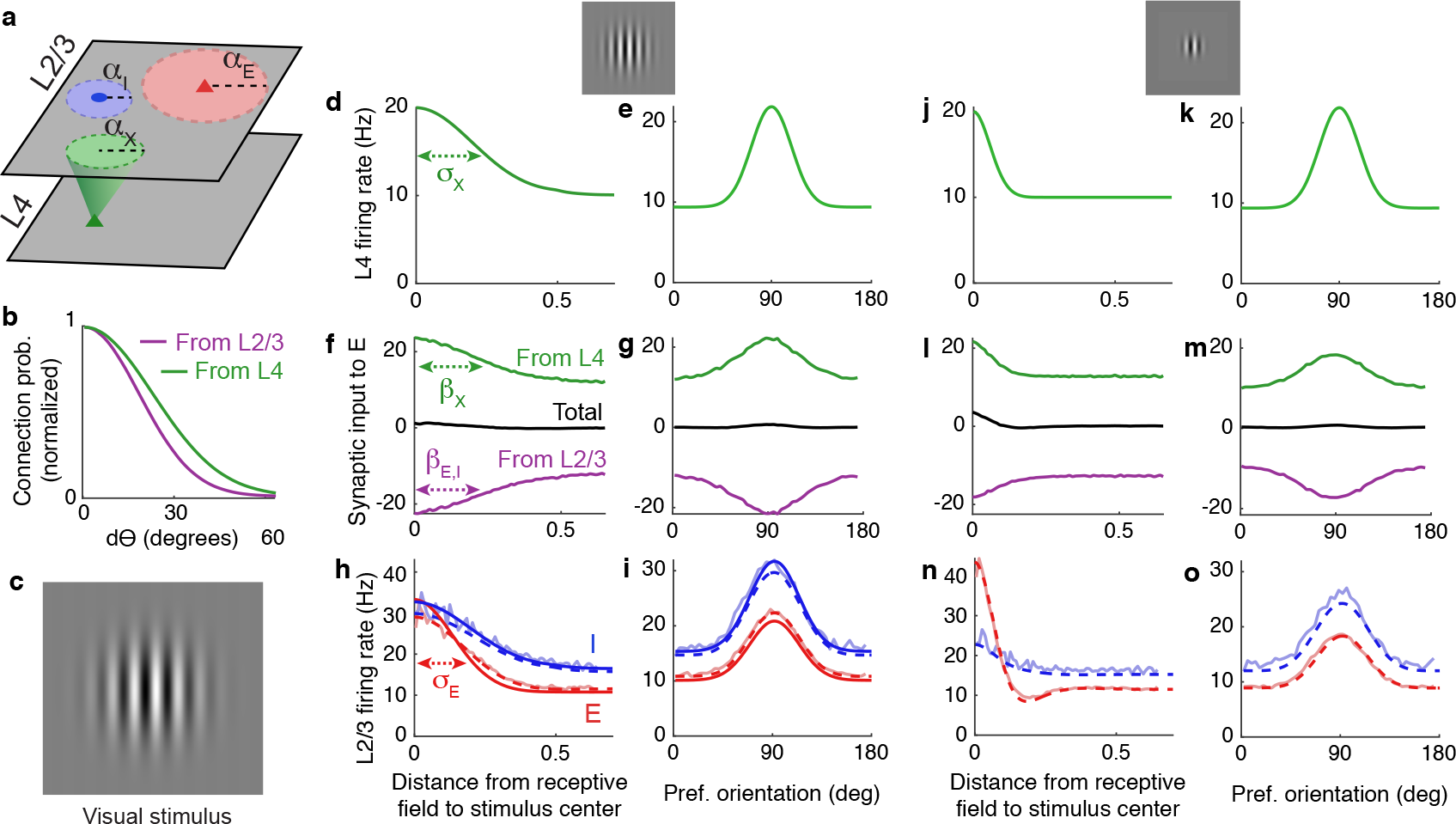
Response properties of a continuously indexed network. **a)** Network diagram. Poisson spiking neurons in L4 (*X*) provide external synaptic input to 2 × 10^5^ recurrently connected excitatory and inhibitory AdEx model neurons (*E* and *I*) in L2/3. The spatial width of synaptic projections from population *a* = *X*, *E*, *I* is given by *β*_*a*_. **b)** Neurons are assigned random orientations and connection probability also depends on the difference, *dθ*, between neurons’ preferred orientation. **c)** An oriented stimulus in the animal’s visual field. **d,e)** The location of the stimulus is modeled by firing rates in L4 that are peaked at the location of the stimulus in physical and orientation space. **f,g)** Synaptic current to neurons in population *E* from the external network (green), the local network (purple) and total (black) as a function distance from the receptive field center and as a function of neurons’ preferred orientation. **h,i)** Firing rate profiles of excitatory (red) and inhibitory (blue) neurons in the local network from simulations (light curves), classical balanced network theory (solid, dark curves; from Eq. (13)) and under the linear correction (dashed; from Eq. (17)) in physical and orientation space. **j-o)** Same as (d-i) except for a smaller visual stimulus, modeled by a narrower spatial firing rate profile in L4.

#### 2.3.1 Amplification and suppression from spatially narrow stimuli

An oriented stimulus localized in the animal’s visual field (Fig. 5c) was modeled by imposing firing rate profiles in L4 that were peaked at the associated location in physical and tuning space, again with a Gaussian profile (Fig. 5d,e). This produced external input to L2/3 that was similarly peaked, but was nearly perfectly canceled by net-inhibitory lateral input (Fig. 5f,g). Excitatory and inhibitory firing rate profiles in L2/3 were also peaked at the associated location in physical and tuning space (Fig. 5h,i), demonstrating that neurons in L2/3 were appropriately tuned to the stimulus.

A smaller visual stimulus was modeled by shrinking the spatial profile of firing rates in L4 while leaving the orientation-dependence of L4 rates unchanged (Fig. 5j,k). As above, synaptic inputs and firing rate profiles were appropriately peaked in physical and orientation tuning space (Fig. 5l-o). However, the smaller stimulus produced a surprising change to firing rates in L2/3. Despite the fact that L2/3 neurons at all locations received less excitation from L4 (Fig. 5l), peak firing rates in L2/3 increased and a surround suppression dynamic emerged (Fig. 5n). Hence, a more localized external input produced an amplification and suppression dynamic similar to the one observed in our model of optogenetic stimulation (compare to Fig. 2). On the other hand, responses in orientation tuning space were mostly unchanged by the smaller stimulus size (Fig. 5m,o).

#### 2.3.2 Mean-field theory of balance in two-dimensional spatial networks with orientation-tuning-specific connectivity

The mean-field theory of balanced networks was previously extended to continuously indexed networks in one and two dimensions [34, 48, 49]. We now review a straightforward extension to two spatial dimensions and one orientation dimension. Eq. (1) generalizes naturally to

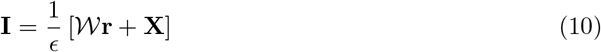

where **r**(**x**, *θ*) = [*r*_*E*_ (*x*, *θ*) *r*_*I*_(*x*, *θ*)]^*T*^ is the vector of mean firing rates of excitatory and inhibitory L2/3 neurons near spatial coordinates **x** = (*x*, *y*) with preferred orientation near *θ*, and similarly for the neurons’ external input, **X**(**x**, *θ*), and total input, **I**(**x**, *θ*). The external input is given by 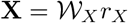 where *r*_*X*_ (**x**, *θ*) is the profile of firing rates in L4 The connectivity kernels, 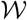 and 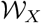, are convolution integral operators defined by

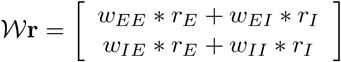

and

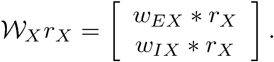

Here, *w*_*ab*_(*x*, *θ*) is the mean-field connection strength between neurons separated by **x** in physical space and *θ* in orientation space (see Methods), and [*w*_*ab*_ ∗ *r*_*b*_](**x**, *θ*) denotes circular convolution with respect to **x** and *θ*, *i.e.*, convolution with periodic boundaries. These convolution operators implement low-pass filters in orientation and physical space, capturing the effects of synaptic divergence and tuning-specific connection probabilities. Similar filters describe feedforward connectivity in artificial convolutional neural networks used for image recognition [31].

Taking *ϵ* → 0 in Eq. (10) shows that that firing rates must satisfy

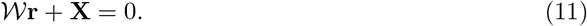

This is an analogue to Eq. (7) for spatial networks. From here, one may be tempted to invert the integral operator 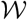 to obtain a spatial analogue of Eq. (3). However, integral operators are never invertible [56]. Specifically, since Eq. (11) is an integral equation of the first kind, there necessarily exist external input profiles, **X**(**x**, 0), for which Eq. (11) does not admit a solution so that the classical balanced state cannot be realized [48]. This implies that there always exist inputs that prevent a continuously indexed network from maintaining excitatory-inhibitory balance. To better understand why this is the case, we follow previous work [5, 34, 48, 50, 49] in transitioning to the spatial Fourier domain to rewrite Eq. (11) as

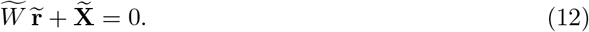

Here, 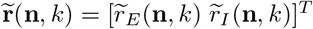 is a Fourier coefficient of **r**(**x**, *θ*) and similarly for 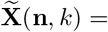 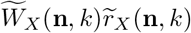 where **n** = (*n*_1_,*n*_2_) is the two-dimensional spatial Fourier mode and *k* is the Fourier mode in tuning space. Importantly, the convolution operators above become ordinary matrices in the Fourier domain. Specifically,

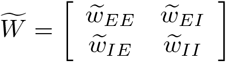

and

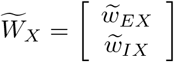

where 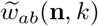 is a Fourier coefficient of *w*_*ab*_(**x**, *θ*). Note that going from Eq. (11) to Eq. (12) requires that 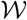 is a convolution operator and that the boundaries of the network are treated periodically, *i.e.*, the convolutions are circular.

Solving Eq. (12) gives an analogue to Eq. (3) for spatial networks in the Fourier domain,

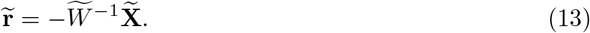

This equation gives all Fourier coefficients, 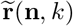. However, this solution is only viable when the inverse transform exists, *i.e.*, when the Fourier series of 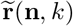 converges, which in turn requires that 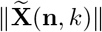 converges to zero faster than 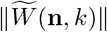 as **n** → 0 and *k* → 0. More specifically, 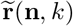 in Eq. (13) must be square-summable. Hence, balance can only be realized when recurrent connectivity, quantified by 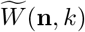, has more power at high spatial frequencies than external input, 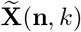. In other words, for balance to be realized, external input, **X**(**x**, *θ*), cannot have “sharper” spatial features than the recurrent connectivity kernels, *w*_*ab*_(**x**, *θ*) for *a, b* = *E, I*.

#### 2.3.3 Balance and imbalance in networks with Gaussian-shaped connectivity kernels

A more intuitive understanding of when and why balance is broken is provided by considering the Gaussian-shaped connectivity and firing rate profiles used in our spiking network simulations. This explanation applies equally to the spatial profile of firing rates and connectivity in physical and orientation space, so we do not distinguish between the two in this discussion. Similar calculations were performed previously for spatial networks [48], so we only review the results here and discuss some of their implications here.

Let *σ*_*a*_ be the width of the Gaussian firing rate profile in population *a*, *α*_*a*_ the width of outgoing synaptic connections from the presynaptic neurons in population *a*, and *β*_*a*_ the width of the spatial profile of synaptic input from population *a* (Fig. 5a,d,f,h). Synaptic divergence broadens the profile of synaptic currents so that

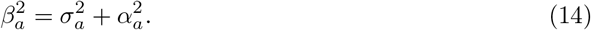

For balance to be maintained, feedforward synaptic input from L4 must be precisely canceled by lateral synaptic input in L2/3. This, in turn, requires that

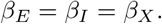

Combined with Eq. (14), this implies that balance requires the widths of firing rate profiles in L2/3 to satisfy [48]

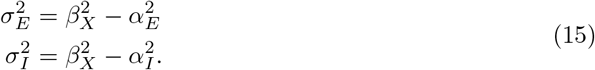

This approximation accurately predicted firing rate profiles in our first spiking network simulation (Fig. 5h,i, solid, dark curves have widths given by Eq. (15)). Hence, by Eq. (15), the requirement of cancellation in balanced networks implies that recurrent connectivity sharpens neurons’ tuning, both in physical and orientation space.

Interestingly, Eq. (15) implies that the amount by which excitatory and inhibitory firing rate profiles are sharpened in balanced networks is determined by the width of their *outgoing* synaptic projections. Pyramidal neurons in L2/3 of mouse V1 preferentially target similarly tuned neurons in L2/3, but the tuning of these lateral connection probabilities is much broader than the tuning of pyramidal neurons’ firing rates [28] (*α*_*E*_ > *σ*_*E*_ in orientation space). This observation is consistent with Eq. (15): Excitatory neuron tuning curves are sharpened precisely because their outgoing connections are broadly tuned. Hence, sharpening of excitatory neuron tuning curves in L2/3 is naturally achieved in balanced networks with lateral excitation, without requiring lateral inhibition. Following the same line of reasoning, the broader orientation tuning of inhibitory neurons [23] (*σ*_*I*_ larger) suggests that they project more locally in orientation tuning space than pyramidal neurons (*α*_*I*_ < *α*_*E*_ in orientation space).

Eqs. (15) also clarify when and why balanced network theory fails for continuously indexed networks. If external inputs are sharper than lateral connectivity (*β*_*X*_ < *α*_*E*_ or *β*_*X*_ < *α*_*I*_) in physical or orientation space, then Eqs. (15) do not yield real solutions for *σ*_*E*_ or *σ*_*I*_. In other words, balance requires that

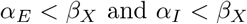

because Eq. (11) does not admit a solution when these inequalities are broken [48]. In other words, the classical balanced state cannot be realized when external synaptic input is too localized for the recurrent network to cancel with its broader connectivity. As a result, balanced network theory cannot be applied to the example in Fig. 5j-o with a smaller visual stimulus.

#### 2.3.4 A linear correction to balance quantifies amplification and suppression in continuously indexed networks

We next derive a linear correction to Eq. (13) that accounts for imperfect cancellation and, in doing so, gives firing rate approximations where classical balanced network theory fails. Specifically, we generalize the derivation of Eq. (4) to continuously indexed networks. Under this linear approximation, firing rate profiles are given by solving

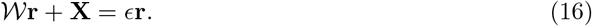

This is an integral equation of the second kind, which generically admits firing rate solutions, **r**, even when Eq. (11) does not [56]. We again transition to the Fourier domain so Eq. (16) becomes

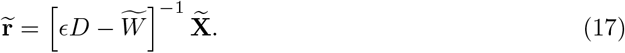

From Eq. (17), firing rates, **r**(**x**, *θ*), can be computed numerically through an inverse transform (the Fourier series over **n** and *k*), yielding an accurate approximation to firing rates from spiking network simulations even where classical balanced network theory fails (Fig. 5n,o).

The amplification and suppression caused by the smaller visual stimulus can be roughly explained by the balanced amplification mechanism discussed previously. Since 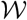 is a low-pass filter, it approximately cancels high frequency components of firing rate profiles. Hence, high frequency components are in the approximate nullspace of the local connectivity operator, 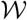, and are therefore amplified by the network through the same mechanism discussed for discrete networks previously.

A more precise explanation is given by first averaging firing rates over orientation preference by setting *k* = 0 in Eq. (17) to give 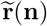 that depends only on spatial frequency, and similarly for 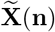 and 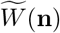. The convolution operator, 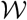, implements a low-pass filter, so 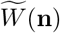 is 𝒪(1) in magnitude at low spatial frequencies and converges to zero at higher frequencies (large ‖**n**‖). The regularized inverse, 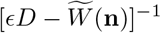, is therefore 𝒪(1) in magnitude at low frequencies and 𝒪(1/*ϵ*) at higher frequencies (Fig. 6a, purple). When external input, **X**(*x*), has sharp features, 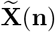 has power at higher spatial frequencies (Fig. 6a, green), which are amplified by the 𝒪(1/*ϵ*) component of 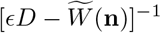 while low frequencies remain 𝒪(1). The result is that the magnitude of 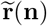 has a 𝒪(1/*ϵ*) peak at a non-zero spatial frequency (Fig. 6a, black), introducing a high-amplitude, non-monotonic rate profile (as in Fig. 5n; see [50] for a similar analysis). When **X**(**x**) has spatially broad features, 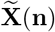 has little power at high spatial frequencies so that this amplification dynamic is weak or absent (as in Fig. 5h). An identical argument applies in orientation space. In summary, high-frequency components of external input profiles are transmitted more strongly than low-frequency components in strongly coupled networks, and the cutoff frequency is determined by the width (*α*_*E*_ or *α*_*I*_) of lateral synaptic projections.

**Figure 6:**
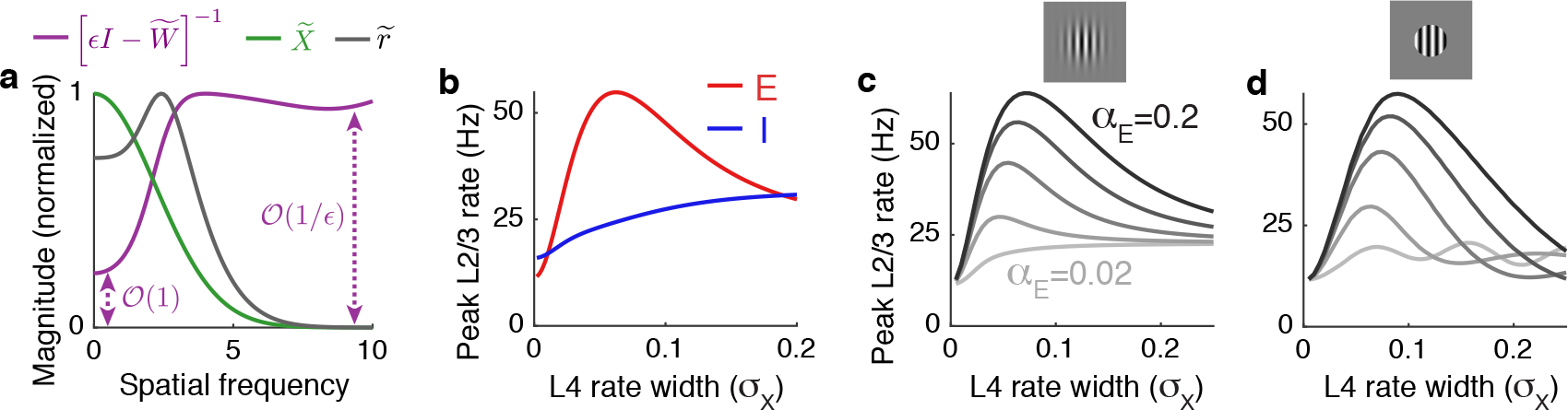
Spatial filtering of external input and the dependence of suppression on outgoing synaptic projection width. **a)** The magnitude of the spatial filter, 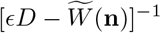, imposed by recurrent connections (purple), the external input (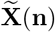, green) and the resulting firing rate profile (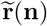, black) as a function of the spatial frequency, 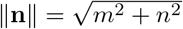, from the simulation in Fig. 5j-o. Magnitude is measured by the Frobenius norm for 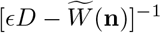. Curves normalized by their peaks. **b)** Firing rates of excitatory (red) and inhibitory (blue) neurons with receptive fields at the center of a grating stimulus plotted as the width of the stimulus increases (represented by increasing *σ*_*X*_) using parameters from Fig. 5j-o. **c)** Same as b, but the excitatory rate is plotted for different widths of the excitatory synaptic projection width, *α*_*E*_. **d)** Same as c, but firing rates in L4 are shaped like a disc with radius *σ*_*X*_ instead of a Guassian with width parameter *σ*_*X*_.

It is worth noting that the average firing rates (over all orientations and spatial positions) are given by the zero Fourier coefficient 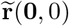. When balance is broken by sharp external input features, the zero Fourier mode is not affected as long as mean firing rates, **r**(**x**, *θ*), remain non-zero at all locations and orientations. Hence, sharp input features can break balance locally without breaking global, network-averaged, balance. This is analogous to the global balance obtained in the optogenetic example when local balance was broken at the level of subpopulations (Fig. 2).

#### 2.3.5 Implications of imbalanced amplification on receptive field tuning

We next considered a study by Adesnik et al. [1] in which drifting grating stimuli of varying sizes were presented to mice while recording from neurons in L2/3 of V1. In that study, pyramidal neurons’ firing rates first increased then decreased as the stimulus size was increased. On the other hand, somatostatin-expressing (SOM) neuron’s firing rates increased monotonically with stimulus size. Intracellular recordings combined with optogenetic stimulation in that study showed that SOM neurons project locally and pyramidal neurons form longer range projections.

To test our model against these findings, we applied Eq. (17) to a network with local inhibition and longer-range excitation (*α*_*E*_ > *α*_*I*_) with increasing size of a visual stimulus (increasing *σ*_*X*_). Our results are consistent with recordings in Adesnik *et al.,* 2012 [1]: Excitatory neuron firing rates changed non-monotonically with stimulus size, while inhibitory neuron firing rates monotonically increased (Fig. 6b). The non-monotonic dependence of excitatory firing rates on stimulus size in Fig. 6b is explained by the mechanism of imbalanced amplification. When *σ*_*X*_ is sufficiently small, balance is broken so imbalanced amplification introduces a large peak firing rate surrounded by suppression (as in Fig. 5n). However, the total amount of external excitation introduced by the stimulus is proportional to the size of the stimulus, so a very small *σ*_*X*_ introduces very little excitation and peak firing rates are small. As *σ*_*X*_ increases, more excitation is recruited and the network is still imbalanced, which leads to increasingly large peak firing rates (as in Fig. 5n). Once *σ*_*X*_ becomes large, balance begins to be restored and the peak excitatory firing rate decreases to moderate values (as in Fig. 5h).

The degree to which excitatory neurons suppress depends on the spatial width, *α*_*E*_, of lateral excitatory projections (Fig. 6c) and suppression of inhibitory neurons similarly depends on the spatial width, *α*_*I*_, of lateral inhibition (not pictured). Specifically, suppression occurs when lateral connectivity is broader than feedforward input (*α*_*E*_ > *β*_*X*_ or *α*_*I*_ > *β*_*X*_) because this is when the balanced solution in Eq. (15) disappears. When a sub-population’s lateral connectivity is more localized than feedforward connectivity from L4 (*α*_*E*_ < *α*_*X*_ as in the lightest gray curve in Fig. 6c; or *α*_*I*_ < *α*_*X*_), that sub-population cannot exhibit suppression since feedforward input width 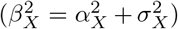 is always larger than lateral connectivity, regardless of the stimulus size (*σ*^*X*^).

A similar line of reasoning explains why peak inhibitory neuron firing rates increase monotonically with stimulus size in Fig. 6b. Inhibitory neurons in that example project locally (*α*_*I*_ = *α*_*X*_), so the inequality *α*_*I*_ < *β*_*X*_ is always satisfied because 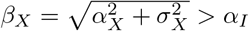 whenever *α*_*I*_ ≤ *α*_*X*_. Whenever *α*_*I*_ < *β*_*x*_, inhibitory firing rates reflect their balanced state values which increase monotonically with the increase in total excitation induced by a larger stimulus.

Unlike SOM neurons, parvalbumin-expressing (PV) neurons were found to exhibit suppression by Adesnik *et al.,* 2012 [1]. Hence, our theory predicts that PV neurons project more broadly in space than SOM neurons. Indeed, PV interneurons in L2/3 are primarily basket cells whose axons project to larger lateral distances than other inhibitory neuron subtypes such as Martinotti cells that comprise most SOM neurons [24].

We observed a unimodal dependence of firing rate on stimulus size (Fig. 6c, all curves have a single peak). However, Rubin, *et al.* [50] observed a multi-modal, oscillatory dependence of firing rate on stimulus size in recordings and in a computational model. In that study, the drifting grating stimuli were disc-shaped with a sharp cutoff of contrast at the edges of the disc. Above, we considered a Gaussian-shaped contrast profile with soft edges (Fig. 6c, inset). Repeating our calculations with a sharp-edged, disc-shaped stimulus (Fig. 6d, inset) produced an oscillatory dependence of firing rate on stimulus size (Fig. 6d), as observed by Rubin *et al.․* This oscillation only arose when lateral synaptic projections were narrower than the stimulus size (*α*_*E*_ small). The oscillation results from a Gibbs phenomenon: The sharp edge in the stimulus produces high-frequency power in 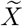, which passes through the high-pass filter 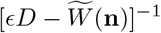 when *α*_*E*_ is small.

We next explored the functional consequences of these results on receptive field tuning. We first considered a disc-shaped grating stimulus (Fig. 7a), producing a disc-shaped firing rate profile in L4 (Fig. 7b). Synaptic divergence causes the profile of synaptic input from L4 to L2/3 to be “blurred” at the edges (Fig. 7c), as quantified by the low-pass filter, 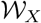. This illustrates a fundamental problem in receptive field tuning: Synaptic divergence from one layer to another implements a low-pass filter that blurs sharp features. This problem is resolved by our observation above that lateral, recurrent connectivity implements a high-pass filter. If the width of lateral, excitatory connections in L2/3 is similar to that of feedforward connections from L4, the high-pass filter implemented by the recurrent network cancels the low-pass filter implemented by feedforward connectivity, effectively implementing a deconvolution that can recover the sharpness of firing rate profiles in L4 (Fig. 7d-f). Hence, counterintuitively, *broader* lateral excitation actually *sharpens* receptive field tuning. Broadening lateral connections further increases the sharpness of the firing rate profiles, but introduce oscillatory, Gibbs phenomena near sharp features (Fig. 7f). These points are illustrated more clearly in an example with an asymmetrically shaped stimulus (Fig. 7g-l). Hence, the high-pass filter described above corrects the blurring caused by synaptic divergence between layers in V1.

**Figure 7:**
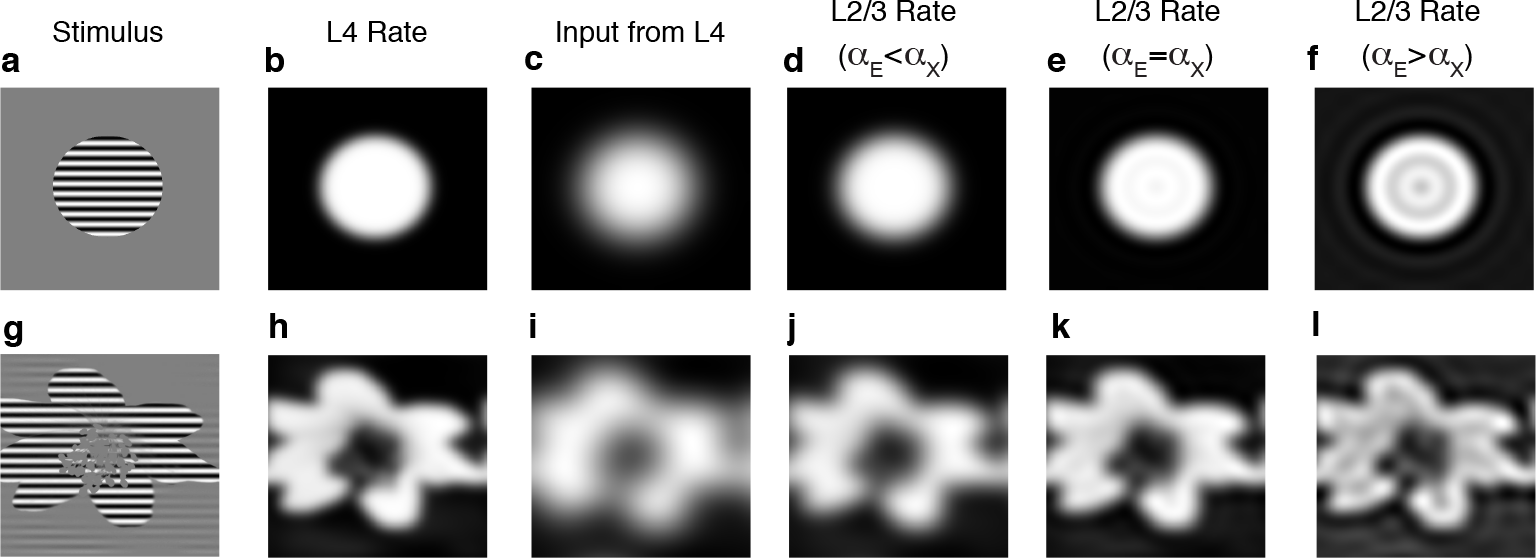
Imbalanced amplification and suppression reverse the blurring introduced by inter-laminar synaptic divergence. **a)** A disc-shaped grating stimulus gives rise to **b)** a disc-shaped firing rate profile, *r*_*X*_(**x**), in L4 with slightly blurred edges (achieved by convolving contrast from a with a Gaussian kernel). **c)** Input, *X*_*E*_(*X*), from L4 to excitatory neurons in L2/3 is blurred by synaptic divergence, which effectively applies a low-pass filter, 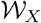, to the L4 rates. **c)** Excitatory firing rates in L2/3 are sharper than external input when lateral excitation is similar, but smaller, in width than interlaminar excitation (*α*_*E*_ = 0.85*α*_*X*_). bf d) Same as c, but lateral excitation is exactly as broad as interlaminar excitation (*α*_*E*_ = *α*_*X*_), which sharpens the edges further, making firing rates in L2/3 similar to those in L4. **d)** Same as c, but lateral excitation is broader than interlaminar excitation (*α*_*E*_ = 1.1*α*_*X*_), which sharpens the edges even further, but also introduced suppressed regions due to Gibbs phenomena. **g-l)** Same as a-f, but contrast was determined by the brightness of a photograph. Horizontal and vertical axes are neurons’ receptive fields.

In summary, imbalanced amplification and linear rate models provide a concise and parsimonious theoretical basis for understanding how suppression, amplification and tuning depends on the profile of neuron’s incoming and outgoing synaptic projections in physical and orientation tuning space.

## 3 Discussion

We described a theory of imbalanced amplification in cortical circuits arising from a local imbalance that occurs when recurrent connectivity structure cannot cancel feedforward input. We showed that imbalanced amplification is evoked by optogenetic stimuli in somatosensory cortex and sensory stimuli in visual cortex, since these stimuli cannot be canceled by the connectivity structure in those areas. Our theoretical analysis of imbalanced amplification explains several observations from cortical recordings in those areas.

Even though firing rates in balanced networks in the large *N* limit do not depend on neurons’ f-I curves (see Eq. (3)), quantifying firing rates under imbalanced amplification relies on a finite size correction that requires an assumption on how firing rates depend on neurons’ input. For simplicity, we used an approximation that assumes populations’ mean firing rates depend linearly on their average input currents, giving rise to Eqs. (4) and (17). In reality, neurons’ firing rates depend nonlinearly on their mean input currents, and also depend on higher moments of their input currents. However, the salient effects of imbalanced amplification are not sensitive to our assumption of linearity. For instance, Eq. (5), which quantifies the strong synaptic currents evoked under imbalanced amplification, does not depend on any assumption about neurons’ f-I curves. The precise value of the firing rates elicited by this strong input does depend on neurons’ f-I curves, however. We found that the linear approximation to f-I curves in Eqs. (4) and (17) performed well at approximating firing rates in our spiking network simulations and also explained several observations from cortical recordings. This may be partly explained due to the fact that our spiking network simulations used neuron models that exhibit spike frequency adaptation, which is known to linearize f-I curves [18, 29] and help networks maintain balance [30]. However, the linear approximation we used cannot explain some phenomena that rely on thresholding and other nonlinear transfer properties [50, 38]. The notion of imbalanced amplification extends naturally to models with nonlinear transfer functions and future work will consider the implications of nonlinearities.

Balanced networks are related to, but distinct from, inhibitory stabilized networks (ISNs) [41, 50, 36] and stabilized supralinear networks that can transition between ISN and non-ISN regimes [50]. The primary distinction is that ISNs are defined by moderately strong recurrent excitation (strong *E* → *E*) whereas balanced networks are defined by very strong external, feedforward excitation (strong *X* → *E*) canceled by similarly strong net-inhibitory recurrent connectivity. Classical balanced networks are necessarily inhibitory stabilized at sufficiently large *N* (small *ϵ*) unless *w*_*EE*_ = 0. However, strongly coupled (approximately balanced) networks can be non-ISN at moderately large *N* (small *ϵ*) if W_*EE*_ is small. Cat V1 is believed to be inhibitory stabilized, which can be used to explain its surround suppression dynamic [41]. However, evidence from optogenetic and electrophysiological studies, suggests that mouse L2/3 V1 might not be inhibitory stabilized: Lateral connection probability is small between pyramidal neurons (small *w*_*EE*_) [24], stimulation of PV neurons does not produce the paradoxical effects that characterize ISNs [4], and modulating pyramidal neuron firing rates only weakly modulates excitatory synaptic currents in local pyramidal neurons [4, 1]. Nonetheless, pyramidal neurons and PV neurons in mouse V1 exhibit surround suppression [1], which we showed is explained by imbalanced amplification.

Despite the similarity in their names, the mechanism of imbalanced amplification studied here is fundamentally different from the mechanism of balanced amplification [39]. First, imbalanced amplification is related to steady-state firing rates, while balanced amplification is a dynamical phenomenon. Moreover, balanced amplification is intrinsic to the local, recurrent circuit: It produces large firing rate transients when local, recurrent inhibition is inefficient at canceling local, recurrent excitation. Imbalanced amplification, on the other hand, produces large steady state firing rates when local, recurrent input is unable to effectively cancel feedforward, external excitation.

The analysis of our spatially extended network model relied on an assumption of periodic boundaries in space, which are not biologically realistic, but approximate networks with more realistic boundary conditions [48]. Without periodic boundary conditions, the integral equations, (10), (11), and (16) are equally valid, but the integrals are defined by regular convolutions in space instead of circular convolutions. As a result, the spatial Fourier modes do not de-couple, so Eqs. (12), (13), and (17) are no longer valid, though they should still offer a good approximation when connectivity is much narrower than the the spatial domain [48]. In addition, anisotropic connectivity statistics, arising for example from tuning dependent connectivity in visual cortical circuits with coherent orientation maps [6], would prevent the integral operator in Eqs. (10), (11), and (16) from being a convolution operator, and therefore preclude the use of Fourier series for the solution. Future work will consider the effects of non-periodic boundaries and non-convolutional connectivity kernels on spatially extended balanced networks.

We focused on firing rates, but sensory coding also depends on variability and correlations in neurons’ spike trains. Our previous work derived the structure of correlated variability in heterogeneous and spatially extended balanced networks when connectivity structure prevents positive and negative correlations from cancelling, effectively providing an analogous theory of imbalanced amplification of correlated variability [49]. Combining those findings with the theory of steady-state firing rates presented here could yield a more complete theory of neural coding in cortical circuits and the effects of imbalanced amplification on coding.

## Acknowledgments

This work was supported by National Science Foundation grants DMS-1517828, DMS-1654268, and DBI-1707400. We thank Ashok Litwin-Kumar for helpful comments on a draft of the manuscript.

## Methods

We modeled recurrently connected networks with *N* neurons, composed of *N*_*E*_ = 0.8*N* excitatory and *N*_*I*_ = 0.2*N* inhibitory neurons. The recurrent network receives external input from a network of *N*_*X*_ neurons that drive the recurrent network. The membrane potential of neuron j from the excitatory (*a* = *E*) or inhibitory (*a* = *I*) population has Adaptive Exponential integrate-and-fire dynamics,

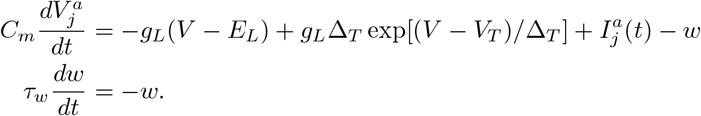

Whenever 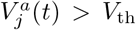, a spike is recorded, the membrane potential is held for a refractory period *τ*_ref_ then reset to a fixed value *V*_re_, and *w* is incremented by *B*. Neuron model parameters for all simulations were *τ*_*m*_ = *C*_*m*_/*g*_*L*_ = 15ms, *E*_*L*_ = —72mV, *V*_*T*_ = —60mV, *V*_th_ = − 15mV, Δ_*T*_ = 1.5mV, *V*_re_ = −72mV, *τ*_ref_ = 1ms, *τ*_*w*_ = 150ms and *B*/*C*_*m*_ = 0.267mV/ms. Membrane potentials were also bounded below by *V*_*lb*_ = −100mV. Synaptic input currents were defined by

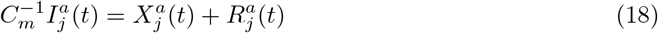

where 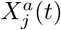 is the feedforward input and 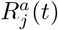 the recurrent input to neuron *j* in population *a* = *E*, *I*. The recurrent input was defined by

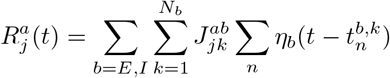

where 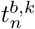 is the nth spike time of neuron k in population *b* = *E, I*. The external input to the recurrent network is defined similarly by

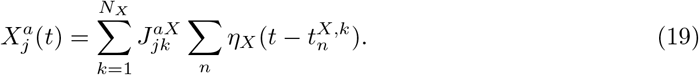

where 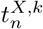 is the nth spike time of neuron *k* = 1,…, *N*_*X*_ in population *X*. Each coefficient, 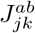, represents the synaptic weight from presynaptic neuron *k* in population *b* to postsynaptic neuron *j* in population *a*. For all simulations, we modeled synaptic kinetics using *η*_*b*_(*t*) = exp(−t/*τ*_*b*_)/*τ*_*b*_ for *t* > 0 where *τ*_*E*_ = 8ms, *τ*_*I*_ = 4ms, and *τ*_*X*_ = 10ms. Note that the integral of *η*_*b*_(*t*) over time is equal to 1 for all three kernels, so the choice of time constant, *τ*_*b*_, does not effect time-averaged synaptic currents. We used *τ*_*I*_ < *τ*_*E*_ < *τ*_*X*_ to prevent excessive synchronous events that break the balanced state. While inhibition may be faster than excitation in many cortical circuits, excitatory neurons are more likely to contact distal dendrites and inhibitory neurons are more likely to contact the soma [27, 22], which could make inhibition functionally faster than excitation. In any case, using fast inhibition is common practice in spiking network simulations with strong or dense connectivity [47, 35, 50, 49, 54] and a complete resolution of this issue is outside the scope of this study.

In Figs. 1, 2 and 3 an extra term, *S* = 2 mV/ms, was added to 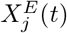 for stimulated neurons during the second half of the simulation to model optogenetic stimulation. We used *N*_*E*_ = 4000, *N*_*I*_ = 1000 and *N*_*X*_ = 4000 (so *N* = 5000) except for Fig. 1f where all *N*_*b*_ values were scaled. Connections were drawn randomly with connection probabilities *p*_*EE*_ = *p*_*IE*_ = *p*_*IX*_ = 0.1, *p*_*EI*_ = *p*_*II*_ = *p*_*EX*_ = 0.2. Since outgoing connections were sampled with replacement, some neurons connected multiple times to other neurons. Synaptic weights were then defined by

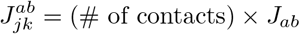

where *J*_*EE*_ = 0.4mV, *J*_*IE*_ = 0.83 mV, *J*_*II*_ = *J*_*EI*_ = -1.67 mV, *J*_*EX*_ = *J*_*IX*_ = 0.47 mV. This gives postsynaptic potential amplitudes between 0.19 and 1.0 mV. For Figs. 1f and 4, the values of *J*_*ab*_ and the values of *p*_*ab*_ were each multiplied by (5000/N)^1/4^ so that they were unchanged at *N* = 5000 and so that 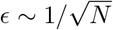. This is slightly different from the more common practice of fixing small connection probabilities and scaling *J*_*ab*_ like 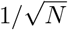. We instead fixed a relatively dense connectivity at *N* = 5000 and the network became increasingly sparse and weakly connected at increased *N*. Both approaches have the same mean-field (since the mean-field only depends on the product of *p*_*ab*_ and *J*_*ab*_), but our approach prevents excessively small synaptic weights at large *N* and prevents dense connectivity at large *N*, which is computationally expensive and susceptible to oscillatory and synchronous spiking.

Spike times in the external population were modeled as independent Poisson processes with **r**_*X*_ =5 Hz. In Fig. 3, external input to the L5 population was created using the spike times of excitatory neurons from the simulations in Fig. 2. Simulations for Fig. 4 were identical to those in Figs. 2 and 3 except there were *N* = 2 × 10^4^ neurons in the L2/3 model, synaptic weights to neurons in that population were multiplied by 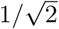, and connections probabilities were also multiplied by 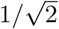. Hence, in relation to Fig. 2, *N* was increased by a factor of four and *ϵ* was halved.

Simulations for Figure 5 used algorithms adapted from previous work [49]. The recurrent network (L2/3) contained *N* = 2 × 10^5^ AdEx model neurons, *N*_*E*_ = 1.6 × 10^5^ of which were excitatory and *N*_*I*_ = 4 × 10^4^ inhibitory. Excitatory and inhibitory neurons in L2/3 were arranged on a uniform grid covering the unit square [0,1] × [0,1] (arbitrary spatial units). The external population (L4) contained *N*_*X*_ = 1.6 × 10^5^ neurons arranged on an identical, parallel square. Each neuron in each population was assigned a preferred orientation chosen randomly and uniformly from 0 to 180°. Connections were chosen randomly as above, but connection probabilities depended on the neurons’ distances in physical and orientation tuning space. Specifically, the connection probability from a neuron in population *b* = *E,I,X* at coordinates **x** = (*x*_1_,*x*_2_) to a neuron in population *a* = *E,I* at coordinates **y** = (*y*_1_, *y*_2_) was

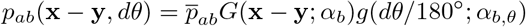

where *dθ* is the difference between neurons’ preferred orientation,

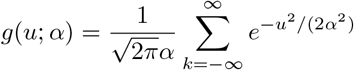

is a one-dimensional wrapped Gaussian and *G*(**u**; *α*) = *g*(*u*_1_; *α*)*g*(*u*_2_; *α*) is a two dimensional wrapped Gaussian. The connection probability averaged over all distances is *β*_*ab*_, which were chosen to be the same as in previous figures, *p*̅_*EE*_ = *p*̅_*IE*_ = *p*̅_*IX*_ =0.1 and *p*̅_*EI*_ = *p*̅_*II*_ = *p*̅_*EX*_ = 0.2. As above, outgoing connections were chosen with replacement, so some neurons made multiple contacts onto other neurons. Connection widths in physical space were *α*_*E*_ = 0.15 and *α*_*I*_ = *α*_*X*_ = 0.04 (as measured on the unit square). Connection widths in orientation space were *α*_*E,θ*_ = *α*_*E,θ*_ = 0.1 and *α*_*X,θ*_ = 0.125 (corresponding to widths of 18° and 22.5° when measured in degrees). Connection strengths, *J*_*ab*_, were the same as in Figs. 1, 2 and 3 except multiplied by a factor of 1.2. Each neuron in L4 was modeled as a Poisson process with rate given by

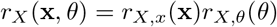

where **x** is the location of the neuron, *θ* is its preferred orientation,

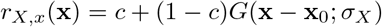

and

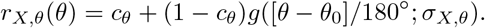

This models a stimulus with orientation *θ*_0_ = 0.5 (representing 90°) and centered at spatial coordinates **x**_0_ = (0.5, 0.5). The parameters *σ*_*X*_ and *σ*_*X,θ*_ quantify the width of L4 firing rates in physical and orientation space. For all panels in Fig. 5, we used *σ*_*X,θ*_ = 0.1 (width 18°) and *c*_*θ*_ = 0.75. We used *σ*_*X*_ = 0.2 for Fig. 5d-i and *σ*_*X*_ = 0.06 for Fig. 5j-o. In both cases, we chose *c* so that the minimum and maximum of *r*_*X,x*_(**x**) were 10 and 20 Hz respectively.

For the spatially extended network, the connectivity kernels, 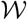 and 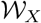, are defined in Results where *w*_*ab*_(**x**,*θ*) = *J*_*ab*_*N*_*b*_*p*_*ab*_(**x**,*θ*)/(*J*_*EX*_*p*_*EX*_*N*_*X*_). The Fourier series in physical and orientation tuning space is defined by

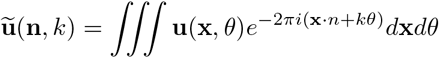

where the triple integral is over the two dimensions of physical space and one dimensional orientation space. The Fourier series of the convolution kernels defined above turns convolution into multiplication in the Fourier domain, from which Eq. (10) gives 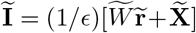 where 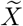, 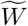, and 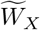 are defined in Results with with 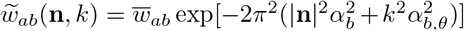, 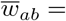 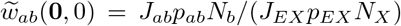, and 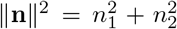. Using the linear approximation, **r** = *g***I** then gives Eq. (17). Firing rates for dashed curves in Fig. 5 and all firing rates in Figs. 6 and 7 were obtained by first computing Eq. (17), then inverting the Fourier transform numerically using an inverse fast Fourier transform. Solid curves in Fig. 5 were computed similarly, except using Eq. (13) in place of Eq. (17).

All simulations and numerical computations were performed on a MacBook Pro running OS X 10.9.5 with a 2.3 GHz Intel Core i7 processor. All simulations were written in a combination of C and Matlab (Matlab R 2015b, MathWorks). The differential equations defining the neuron model were solved using a forward Euler method with time step 0.1 ms.

